# The landscape of the histone-organized chromatin of Bdellovibrionota bacteria

**DOI:** 10.1101/2023.10.30.564843

**Authors:** Georgi K. Marinov, Benjamin Doughty, Anshul Kundaje, William J. Greenleaf

**Affiliations:** Department of Genetics, Stanford University, Stanford, California 94305, USA; Department of Computer Science, Stanford University, Stanford, California 94305, USA; Center for Personal Dynamic Regulomes, Stanford University, Stanford, California 94305, USA; Department of Applied Physics, Stanford University, Stanford, California 94305, USA; Arc Institute, Palo Alto, California, USA

## Abstract

Histone proteins have traditionally been thought to be restricted to eukaryotes and most archaea, with eukaryotic nucleosomal histones deriving from their archaeal ancestors. In contrast, bacteria lack histones as a rule. However, histone proteins have recently been identified in a few bacterial clades, most notably the phylum Bdellovibrionota, and these histones have been proposed to exhibit a range of divergent features compared to histones in archaea and eukaryotes. However, no functional genomic studies of the properties of Bdellovibrionota chromatin have been carried out. In this work, we map the landscape of chromatin accessibility, active transcription and three-dimensional genome organization in a member of Bdellovibrionota (a *Bacteriovorax* strain). We find that, similar to what is observed in some archaea and in eukaryotes with compact genomes such as yeast, *Bacteriovorax* chromatin is characterized by preferential accessibility around promoter regions. Similar to eukaryotes, chromatin accessibility in *Bacteriovorax* positively correlates with gene expression. Mapping active transcription through single-strand DNA (ssDNA) profiling revealed that unlike in yeast, but similar to the state of mammalian and fly promoters, *Bacteriovorax* promoters exhibit very strong polymerase pausing. Finally, similar to that of other bacteria without histones, the *Bacteriovorax* genome exists in a three-dimensional (3D) configuration organized by the parABS system along the axis defined by replication origin and termination regions. These results provide a foundation for understanding the chromatin biology of the unique Bdellovibrionota bacteria and the functional diversity in chromatin organization across the tree of life.

## Introduction

Histones and nucleosomal chromatin are one of the defining features of the broadest divisions of life on our planet. Nearly all eukaryotes ^6^ package their chromatin around nucleosomal particles, composed of two dimerized tetramers of the four core histones H2A, H2B, H3 and H4, and histones are the most conserved proteins in their genomes ^7^, especially when it comes to the key residues playing a vital role in the so called “histone code” ^8^ (and through it in all aspects of chromatin biology – transcription, gene regulation, DNA repair, DNA replication, mitosis and meiosis, and many others), which are almost universally invariant across all eukaryotes.

Eukaryotic histones trace their ancestry to Archaea ^9^, which is the half of the prokaryote diversity from within which eukaryotes appear to have emerged, and where the informational processing systems of eukaryotes are thought to derive from ^10–15^. However, archaeal histones are quite different from those of eukaryotes. While they all contain the same canonical histone fold domain ^16^, consisting of three alpha helices ^17^ dimerizing with another histone molecule, archaeal histones only contain the histone fold domain ^18,19^ while those of eukaryotes have long tails (that are the primary sites of post-translational modifications). Another major difference is that unlike the eukaryotic octamer nucleosome, archaeal histone dimers can oligomerize into so called “hypernucleosomes” ^20–23^, which are stacks of individual histone dimers, and can be of variable length.

While bacteria (the other main branch of prokaryotes) do not have histones as a rule, recent phylogenomic and experimental studies have revealed that some bacteria in fact do possess histone proteins ^24,25^. Most notably in the bacterium *Bdellovibrio bacteriovorus* and other members of the Bdellovibrionota phylum, histones are both abundant and essential for viability components of the nucleoid ^26,27^. Bdellovibrionota histones have been shown to bend DNA as dimers and in a sequence-independent manner ^27^, while other studies have suggested that they exhibit yet another major deviation from the conventional eukaryote state of nucleosomal architecture by binding to DNA head-on to coat it, as opposed to binding to DNA by wrapping the DNA strand around histones proteins ^26^.

These observations pose a long list of tantalizing questions about the physical organization of the genomes of these histone-possessing bacteria, about the relationship between chromatin structure and gene expression, and about how such features compare to those of archaea. However, while archaea have received some experimental attention in the past decade ^28–31^, these properties have not been assayed so far with functional genomic tools in the bacterial clades that have histones.

To begin addressing these outstanding issues, we have mapped chromatin accessibility using ATAC-seq ^32^ (Assay for Transposase-Accessible Chromatin using sequencing),transcriptional activity using KAS-seq ^35^ (Kethoxal-assisted single-stranded DNA sequencing), and three-dimensional genome organization using Hi-C ^36^ in a member of the Bdellovibrionota phylum (a *Bacteriovorax* sp. ICPB 3264 [H-I A3.12] strain). Our data reveals a chromatin accessibility landscape reminiscent of that of eukaryotes with highly compact genomes, with preferentially accessible promoter regions, strong polymerase pausing at promoters, positive correlation between promoter accessibility and active transcription rates, and three-dimensional organization similar to that of other bacteria. We discuss these properties in the broad context of the diversity of life on our planet.

## Results

### Genome assembly and annotation of *Bacteriovorax* sp. ICPB 3264 [H-I A3.12]

We first set out to map the chromatin accessibility landscape in *Bdellovibrio bacteriovorus* using ATAC-seq, given that *Bdellovibrio* histones were recently structurally characterized ^26^ and *Bdellovibrio bacteriovorus* is the most well-studied Bdellovibrionota representative. However, a major technical challenge to the application of ATAC-seq to *Bdellovibrio* is presented by the unique lifestyle of these bacteria. *Bdellovibrio* is the most famous example of a predatory prokaryote ^37–39^ – it feeds by attaching onto the cell wall of gram-negative bacteria, creating an opening into it, then inserting itself into the periplasmic space between the inner and outer bacterial membranes, breaking down the host cell, subsequently undergoing polyploid division, and eventually lysing the remnants of the host. This means that *Bdellovibrio bacteriovorus* is usually grown together with another prey species, which are bacteria without histones.

This predatory lifestyle makes applying ATAC-seq to *Bdellovibrio* challenging as the basic principle behind ATAC-seq is the strong preference of the Tn5 transposase for open chromatin not protected by nucleosomes ^32^. In mammalian cells, this can result in extreme overrepresentation of mitochondrial DNA sequences in final libraries if mitochondria have not been properly filtered out during the nuclei isolation procedure. In a mixture of bacteria without and with histones, if the latter’s histones are strongly protecting DNA from Tn5 insertion, ATAC-seq libraries could easily consist almost entirely of fragments from the prey rather than *Bdellovibrio*.

Fortunately, prey-independent strains of *Bdellovibrio* are also available, allowing axenic growth in media free of other bacteria. We obtained “*Bdellovibrio bacteriovorus*” strain ICPB 3264 [H-I A3.12] from ATCC and carried out ATAC-seq on it. However, almost no reads from these libraries aligned to any of the previously sequenced Bdellovibrionota strains available in the NCBI databases (neither *Bdellovibrio* nor strains from *Bacteriovorax*, which is the other major division of the phylum).

We thus carried out *de novo* genome sequencing of the strain we worked with, using a combination of nanopore long reads and short Illumina reads (see the Methods section for details). This resulted in a single contig of length 4,148,738 bp (∼667 × coverage) predicted to encode (see Methods for details) 4,127 protein coding genes (Figure 1A).

**Figure 1:**
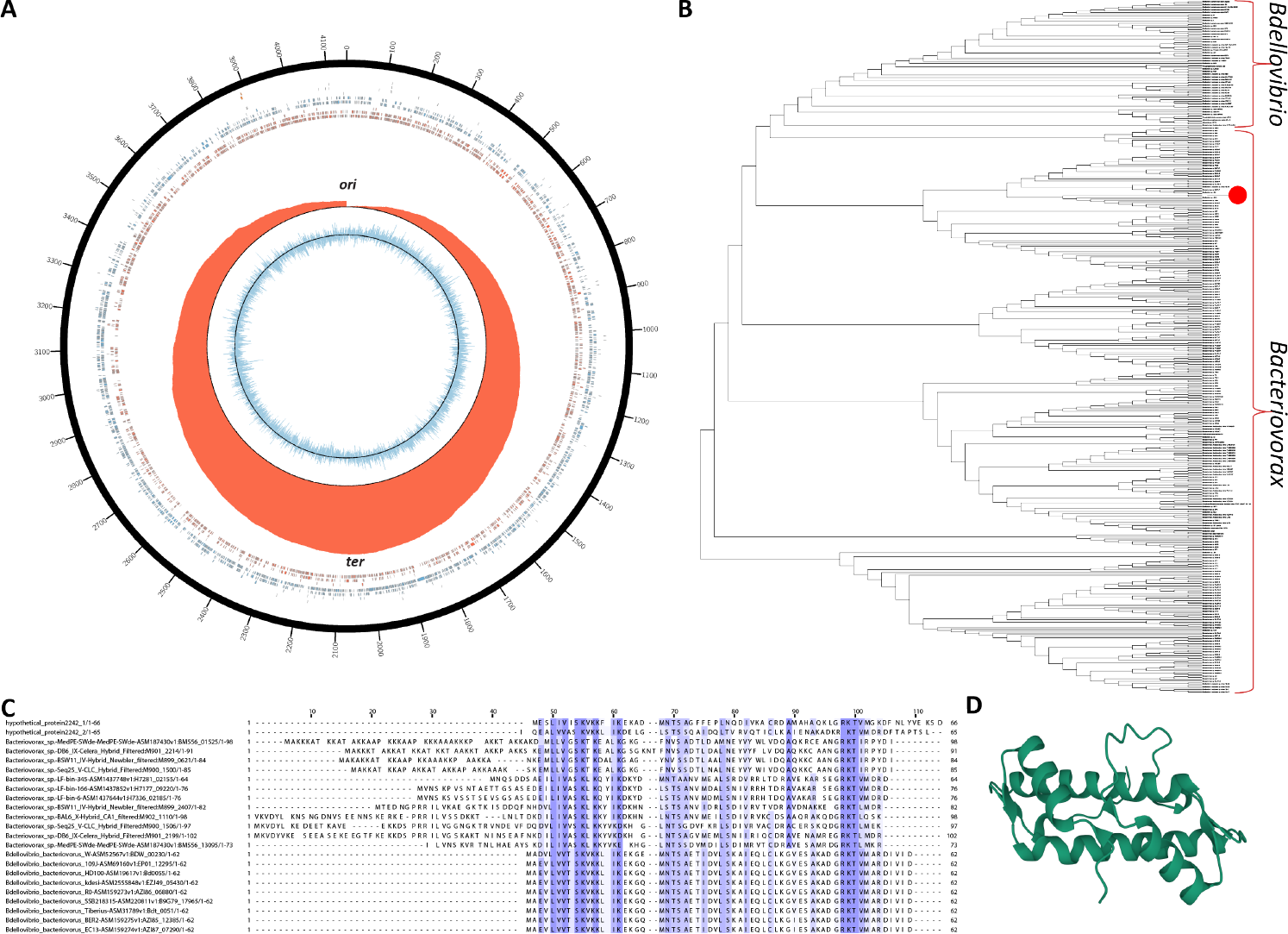
Genome assembly and annotation of *Bacteriovorax* sp. ICPB 3264 [H-I A3.12]. (A) Circos ^1^ plot of the *Bacteriovorax* sp. ICPB 3264 [H-I A3.12] chromosome. Forward- and reverse-strand protein coding genes are shown in red and blue respectively in the inner tile layer; ribosomal RNAs and tRNAs are shown in the outer tile layers. The red histogram shows the cumulative GC-skew ^2^; with the inflection points corresponding to the replication origins and and termini ^3^. (B) Maximum likelihood 16S rRNA tree (generated using RAxML-NG ^4^) of members of the Bdellovibrionota phylum and the phylogenetic location of strain ICPB 3264 [H-I A3.12]. (C) Multiple sequence alignment of histone proteins identified in select *Bacteriovorax* and *Bdellovibrio* genomes. In contrast to the singlet histone folds in *Bdellovibrio*, the histone gene in *Bacteriovorax* sp. ICPB 3264 [H-I A3.12] is a histone doublet; the two histone folds are shown separately (top two rows). (D) Predicted ^5^ protein structure of the histone gene in *Bacteriovorax* sp. ICPB 3264 [H-I A3.12].

Analysis of potential DNA modifications revealed the existence of 5-methylcytosine methylation in a CpGGCC context. This signature is likely associated with a restriction modification (R-M) system ^40^ in *Bacteriovorax*.

We then established the precise phylogenetic positioning of the strain by analyzing available 16S rRNA sequences for Bdellovibrionota strains. The phylum Bdellovibrionota ^41^ includes three major clades/classes – Bacteriovoracia, Bdellovibrionia, and Oligoflexia. Bacteriovoracia includes the *Bacteriovorax, Halobacteriovorax* and *Peredibacter* genera while Bdellovibrionia features *Bdellovibrio* and *Pseudobdellovibrio*. Our phylogenetic analysis points to strain ICPB 3264 [H-I A3.12] beloning to the *Bacteriovorax* genus, being most closely related to *Bacteriovorax* sp. B1T4-F (Figure 1B), even though it was originally labeled as a “*Bdellovibrio*” strain. We refer to it as *Bacteriovorax* sp. ICPB 3264 [H-I A3.12] or simply *Bacteriovorax* in the subsequent text.

Finally, we examined putative *Bacteriovorax* sp. ICPB 3264 [H-I A3.12] histone genes in order to confirm their presence in its genome, and found one histone protein to be encoded in our assembly. Unlike *Bdellovibrio* histone genes, which encode histone singlets that later form dimers, in *Bacteriovorax* sp. ICPB 3264 [H-I A3.12] the histone gene encodes a histone doublet that presumably functions as a dimer on its own (Figure 1C–D).

### ATAC-seq reveals the accessible chromatin landscape in *Bacteriovorax*

In order to map the chromatin accessibility landscape in *Bacteriovorax*, we adapted the previously described bacterial ATAC (bac-ATAC) protocol ^42^, which involves crosslinking with 1% formaldehyde and permeabilization of the cell wall with lysozyme (see the Methods section for details).

In metazoans, ATAC-seq libraries display a clear nucleosomal signature, with a strong subnucleosomal peak (≤ 120 bp) followed by a robust mononucleosomal (∼150 bp) peak, a weaker dinucleosomal peak, and so on. In *Bacteriovorax*, we do not observe a nucleosomal protection pattern but only a single broad fragment peak centered between 100 bp and 200 bp (Figure 2B). Thus *Bacteriovorax* histones do not appear to confer the same arrayed local protection to DNA as eukaryotic ones.

**Figure 2:**
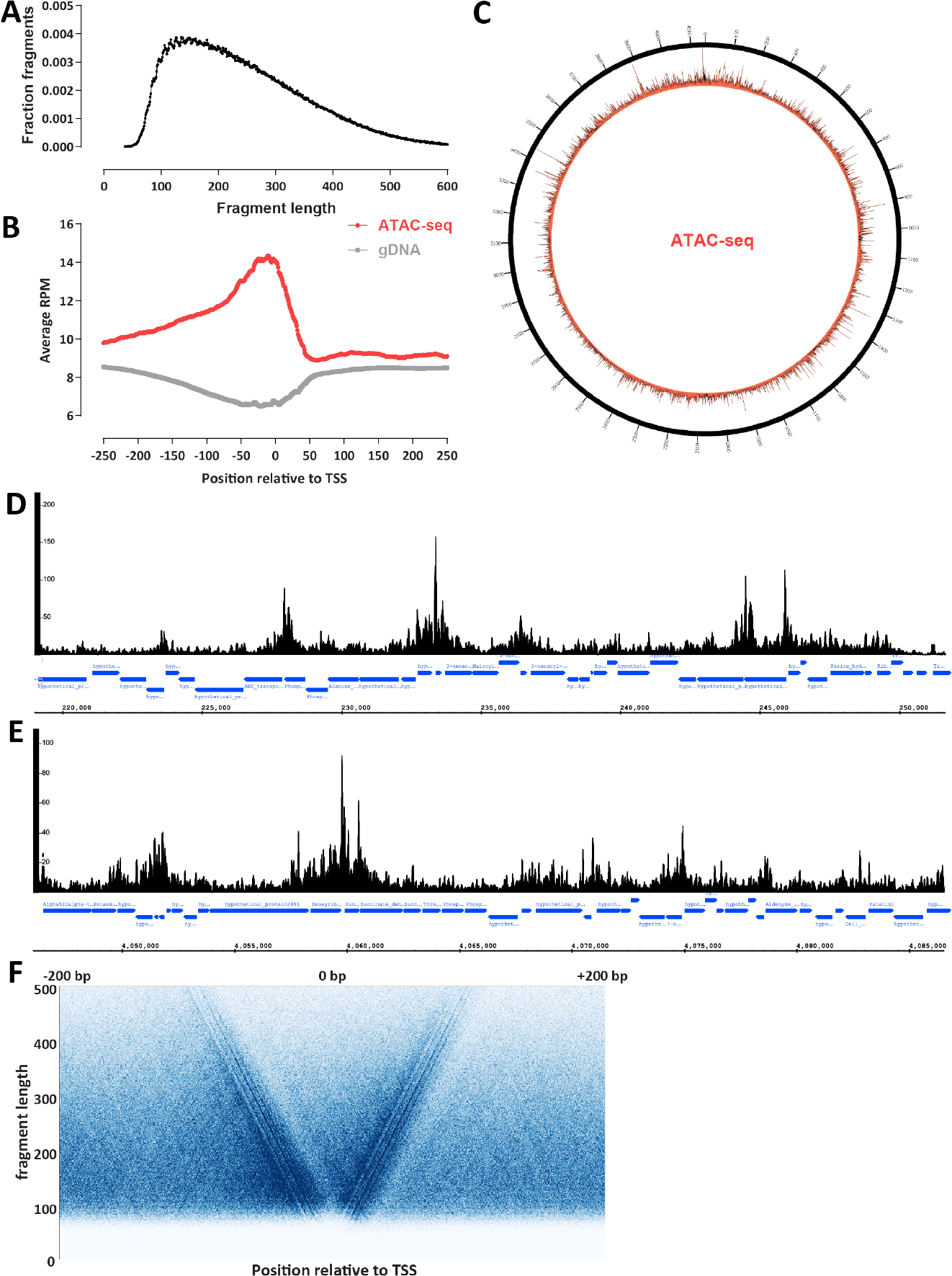
The accessible chromatin landscape in *Bacteriovorax*. (A) ATAC-seq fragment length distribution. (B) ATAC-seq metaprofile around annotated *Bacteriovorax* sp. ICPB 3264 [H-I A3.12] TSSs. (C) Circos plot of the global accessibility distribution along the *Bacteriovorax* sp. ICPB 3264 [H-I A3.12] chromosome. (D-E) Representative snapshots of local ATAC-seq signal distribution in *Bacteriovorax* (exponentially growing culture). (F) V-plot of ATAC-seq fragment distribution around annotated *Bacteriovorax* sp. ICPB 3264 [H-I A3.12] TSSs.

Accessibility is centered around the promoter regions between genes (Figure 2B-E). We observe transcription start sites (TSS) scores (indicating the level of enrichment over promoter regions relative to flanking regions; see the Methods section and previous detailed discussions ^43^) in the 1.35-1.40 neighborhood, which is comparable to what we had previously found in the archaeon *Haloferax volcanii* ^31^ and also to the lower end what is usually seen in eukaryotes with highly compact genomes such as yeast ^44^. In contrast, naked genomic DNA controls exhibit slight depletion around TSS (Figure 2B). Manual inspection of ATAC-seq profiles along the genome confirms these global observations, but also shows frequent cases of elevated accessibility extending into gene bodies (Figure 2D-E).

We also note that a previous ATAC-seq study of conventional bacteria that do not possess histones (*Caulobacter crescentus*) reported the existence of broad domains of elevated and decreased accessibility spanning hundreds of kilobases ^42^. This is not readily observed in *Bacteriovorax* (Figure 2C), just as it is not observed in the archaeon *Haloferax volcanii*.

Finally, we asked whether patterns of nucleosomal positioning are observed around promoters using V-plot analysis ^45^. With the caveat that we do not yet have precise TSS annotations for *Bacteriovorax*, we do not observe strongly positioned nucleosomes around its TSSs (Figure 2F).

### The ssDNA and active transcription landscape in *Bacteriovorax*

Next, we mapped the landscape of active transcription in *Bacteriovorax* using KAS-seq ^35^, which strongly and specifically enriches for ssDNA structures by labeling exposed guanine (G) bases with N_3_-kethoxal, followed by biotin labeling through a click reaction, fragmentation of DNA, and streptavidin pulldown (see the Methods section for details). While KAS-seq labels all ssDNA structures (G-quadruplexes, replication intermediates, etc.), the most abundant source of ssDNA in the cell are transcriptional bubbles associated with RNA polymerases directly engaged with DNA, both paused and actively elongating ^35^.

As with ATAC-seq, we do not observe broad large-scale domains of active transcription along the *Bacteriovorax* chromosome in our KAS-seq datasets (Figure 3A). The global landscape is largely uniform, punctuated by strong localized peaks.

**Figure 3:**
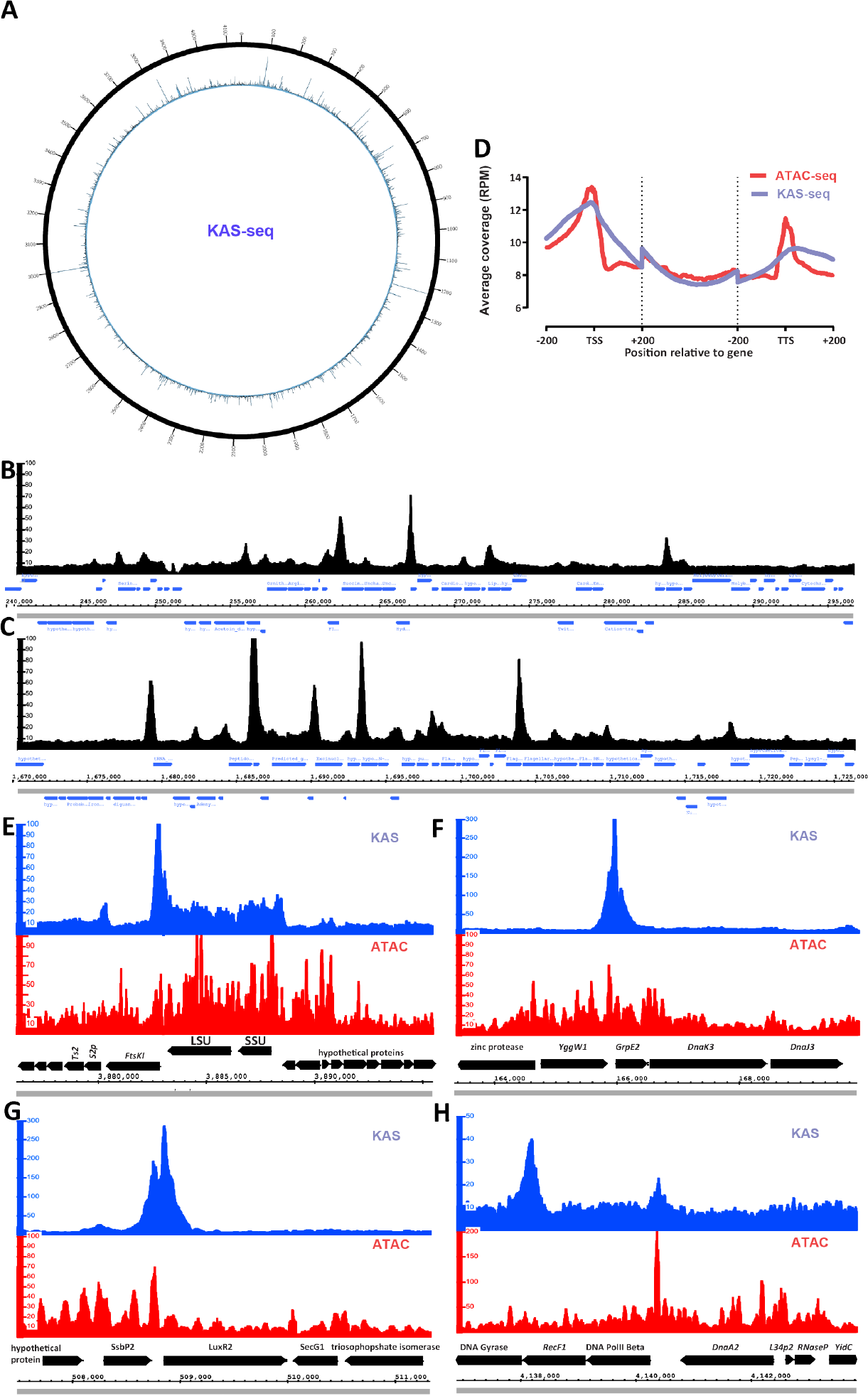
The ss-DNA and active transcription land-scape in *Bacteriovorax*. (A) Circos plot of the global KAS-seq signal distribution along the *Bacteriovorax* sp. ICPB 3264 [H-I A3.12] chromosome. (B-C) Local KAS-seq signal distribution in *Bacteriovorax* shows strongly localized peaks associated with promoter regions (D) Metaplot of KAS-seq and ATAC-seq signal distribution along *Bacteriovorax* genes. (E) KAS-seq and ATAC-seq profiles around the ribosomal RNA locus of *Bacteriovorax*. (F) KAS-seq and ATAC-seq profiles around the heat shock protein *GrpE2* gene. (G) KAS-seq and ATAC-seq profiles around the *LuxR2* transcriptional activator gene. (H) KAS-seq and ATAC-seq profiles near the origin of replication of the *Bacteriovorax* chromosome.

Strikingly, examination of local browser tracks (Figure 3B-C) revealed these peaks to be associated with promoters, indicating the existence of very strong promoter pausing in *Bacteriovorax*. Polymerase pausing and its later controlled release are decisive steps in gene regulation in many eukaryotes ^46^, having been first described in the fruit fly *Drosophila melanogaster* ^47–50^, and also shown to be widespread in mammals ^51,52^, most metazoans, and some plants, but curiously absent in yeast and *Arabidopsis thaliana* ^53^. Bacteria do exhibit sequence-dependent polymerase pausing but this has been primarily reported in the context of transcription elongation and termination ^54–59^ and *E. coli* promoters are not characterized by the typical peaks associated with pausing in global run-on GRO-seq datasets ^53^. The strong KAS-seq peaks we observe are not due to transcription termination-associated pausing because they are found in all possible orientations of gene pairs, including between two divergent promoters (Figure 3B-C).

We had previously observed promoter pausing using KAS-seq in the euryarchaeon *Haloferax volcanii* ^31^. *Bacteriovorax* promoters are even more accentuated in KAS-seq profiles than in *Haloferax* – promoter KAS signal peaks at ∼12 RPM (Reads Per Million mapped reads) over *Bacteriovorax* promoters compared to a through at ∼10 RPM in intergenic space and ∼8 over gene bodies (Figure 3B-D); in *Haloferax* these values are ∼11, ∼10 and ∼8 RPM, respectively ^31^.

The most strongly transcribed (as assessed by KAS-seq levels over the gene body) genes in *Bacteriovorax* are the two ribosomal RNAs (Figure 3E). Notably, these do not exhibit a KAS-seq peak in their promoter (although a very strong such peak is located downstream of the ribosomal RNA operon). This region is also one of the most accessible over the gene body loci in the genome as measured by ATAC-seq; this points to an analogous situation to the one well known from yeast and other eukaryotes. Yeast rDNA exists in two states – a transcriptionally inactive chromatinized condition on one hand, and a highly transcriptionally active, almost entirely devoid of histones state ^44,60,61^ on the other. These observations also contrast with those from ATAC-seq for *Caulobacter crescentus*, which does not possess histones, and in whose genome ribosomal RNA clusters were reported to be one of a handful of highly transposase-inaccessible transcribed regions ^42^ (HINTs).

Other genes where particularly strong putative paused RNA polymerase peaks are observed include the *GrpE2* heat shock protein (Figure 3F), a *LuxR2* transcriptional activator gene (Figure 3G), the chaperone protein ClpB ATP-dependent unfoldase (Supplementary Figure 1A), the RNA Polymerase Sigma Factor RpoH1 (Supplementary Figure 1B), and others. For each of these proteins, especially the heat shock ones, it is plausible that polymerase pausing is a regulatory mechanism specifically designed to enable their very rapid activation, in a manner analogous to heat shock genes in *Drosophila* where the phenomenon was originally described ^47,48^.

The strongest ATAC-seq peak in the genome is found in an unusually large intergenic space without any annotated genes located between replication-related genes and near the origin of replication (Figure 3H). This locus exhibits only a modest ssDNA peak, and may be associated with replication initiation.

### Chromatin accessibility correlates with transcriptional activity in *Bacteriovorax*

Our next step was to characterize the relationship between chromatin accessibility and transcriptional activity in *Bacteriovorax* as well as their dynamics upon large-scale gene expression perturbations. While genome-wide gene expression dynamics in these organisms has not been studied extensively previously, it has been reported that starvation of *Bdellovibrio* cells for 4 hours in HEPES buffer results in major changes in gene expression ^62^.

We carried out ATAC-seq and KAS-seq in multiple replicates from the same cells and at the same time in both exponentially growing and HEPES-starved cultures, and indeed observed large-scale changes in transcriptional activity and chromatin accessibility (Figure 4A-C). Curiously, chromatin accessibility around promoters largely disappeared in starved cells, even though transcriptional activity was either unaffected at many promoters or shifted markedly in others. The overall properties of the active transcription landscape were similar to those of exponentially growing cells, except for a slight shift from promoters towards presumed elongation over gene bodies for a number of genes (Figure 4C). We identified 555/378 gene bodies and 488/607 TSSs with respectively statistically decreased/increased KAS-seq signal.

**Figure 4:**
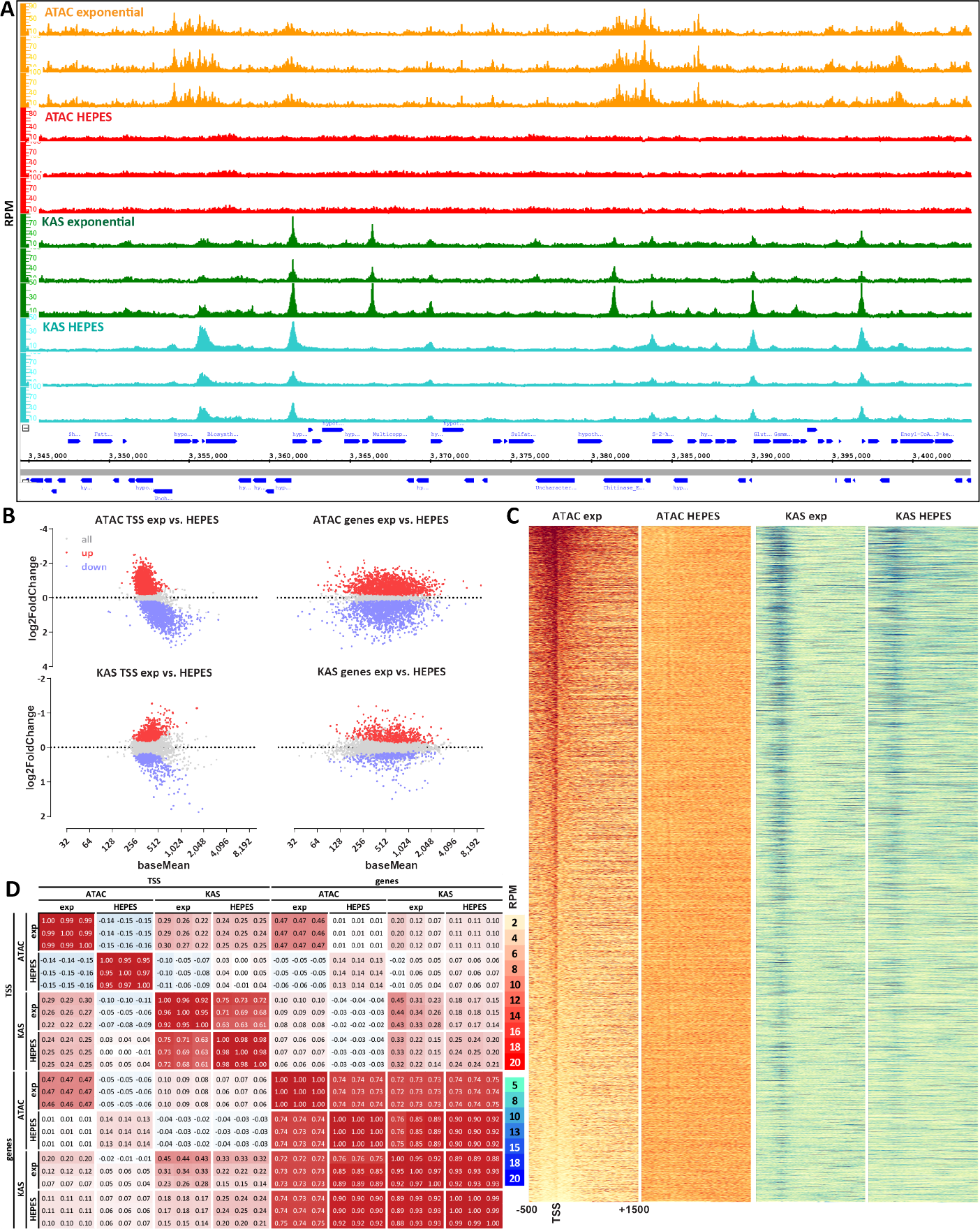
Relationship between chromatin accessibility and transcriptional activity in *Bacteriovorax*. (A) Starvation of *Bacteriovorax* cells induces changes in the chromatin accessibility and active transcription landscapes. Show are three independent KAS-seq and ATAC-seq replicates for each condition. (B) Differential accessibility and KAS levels between exponentially growing and starved (HEPES) cells. (C) Global ATAC-seq and KAS-seq profiles over *Bacteriovorax* genes under exponentially growing and starvation conditions. (D) Correlation between ATAC-seq and KAS-seq signal over TSSs and gene bodies.

Previously we found no correlation between chromatin accessibility and transcriptional activity in the archaeon *Haloferax volcanii* ^31^. In contrast, ATAC-seq signal both over promoter regions and gene bodies correlates positively with KAS signal, again over both promoter regions and gene bodies (Figure 4D).

### Frequent independent transcription initiation/regulation of individual genes within *Bacteriovorax* operons

Functionally related genes are often organized into operons in both bacteria and archaea ^63^ that are transcribed together. However, evidence has accumulated over the years that certain genes within operons may be transcribed and regulated separately from the whole operon ^64–67^. KAS-seq and ATAC-seq data for the archaeon *Haloferax* also demonstrated this phenomenon on a genome-wide scale – individual promoters, discernible by strong ATAC-seq and/or KAS-seq peaks inside operons, are frequently observed in that organism, as are obviously different KAS-seq levels over different genes within a single operon ^31^.

We aimed to determine whether similar unexpected complexity in the functional organization of operons also exists in the genomes of bacteria with histone-based chromatin. To this end we identified cases of clear operons (i.e. colinear genes obviously belonging to the same functional group) and examined KAS-seq and ATAC-seq profiles in both conditions that we assayed (Figure 5). We observed multiple cases of both internal KAS-seq and ATAC-seq peaks inside operons (e.g. Figure 5B and Figure 5E). While internal KAS-seq peaks could be associated with polymerase pausing due to regulation of transcriptional elongation, RNA processing or cotranscriptional translation events, internal ATAC-seq peaks most likely correspond to independent promoters.

**Figure 5:**
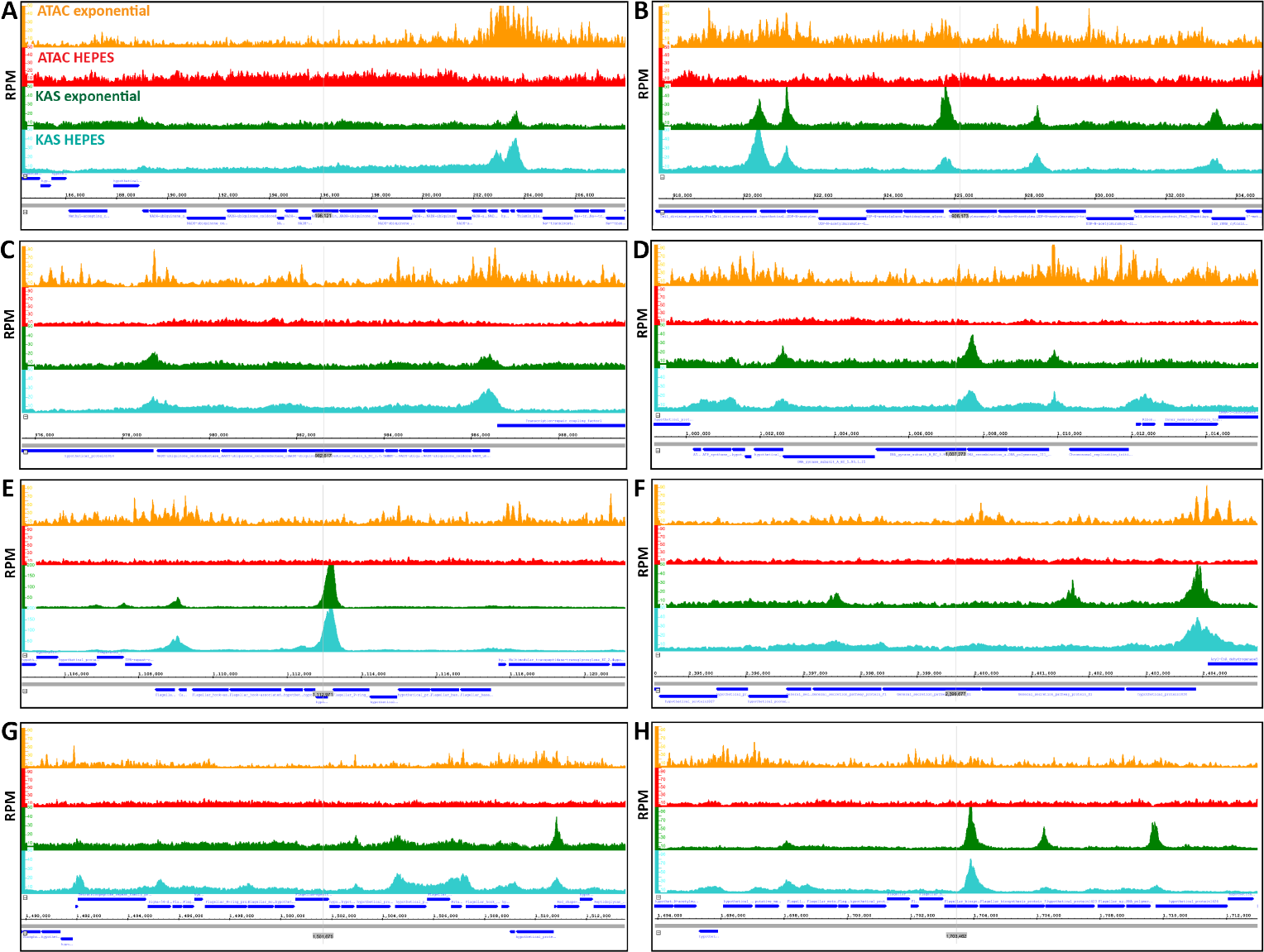
Chromatin accessibility and active transcription levels in *Bacteriovorax* operons. (A-H) Genome browser snapshots of ATAC-seq and KAS-seq levels in exponentially growing and starved (HEPES) cells for eight different *Bacteriovorax* operons.

### Three-dimensional organization of the *Bacteriovorax* genome

Finally, we characterized the 3D organization of the *Bacteriovorax* genome. We employed a modification of the Hi-C ^36^ assay, using multiple restriction 4-cutter enzymes to digest crosslinked *Bacteriovorax* chromatin to improve resolution in its very small and compact genome (see the Methods section for details).

Hi-C has been previously applied to study the physical organization of several bacterial and archaeal strains, such as originally *Caulobacter crescentus* ^68,69^ and later multiple others ^70–77^. These studies have revealed two main typical features of bacterial chromosomes. In *Caulobacter* and most other bacteria, Hi-C maps exhibit a prominent pattern of increased interactions perpendicular to the main diagonal and connecting the origin and the terminus of replication, reflecting a chromosome configuration driven by SMC/condensin complexes loaded at centromere-like parS sites near replication origins ^78^. The parABS system ^79–82^ plays a major role in this process, with most bacteria having one or multiple highly conserved ^83^ parS sites near their origin of replication. The second common major feature are self-interacting chromosomal interaction domains (CIDs), areas of increased local contact frequency corresponding to plectonemes arising due to transcription-induced supercoiling ^68^.

Hi-C maps of archaea differ considerably – *Sulfolobus*, which does not have histones, appears to possess three distinct origins of replication resulting in a much more complicated chromosomal configuration, and does not display CIDs ^84^, while *Haloferax*, which has histones (although it is not clear to what extent they package the genome), was reported to exhibit CIDs but not the perpendicular to the main diagonal feature typical to bacteria ^85^.

In contrast, *Bacteriovorax*, although it does possess histones, exhibits a typical large-scale bacterial organization of its chromosome, with very strong cross-diagonal interactions connecting the replication origin and termination regions (Figure 6A). However, we do not observe clear evidence for plectonemic CID-like structures in our maps. HEPES-starved cells exhibit decreased strength of the cross-diagonal feature (Figure 6B), likely reflecting much lower replication activity.

**Figure 6:**
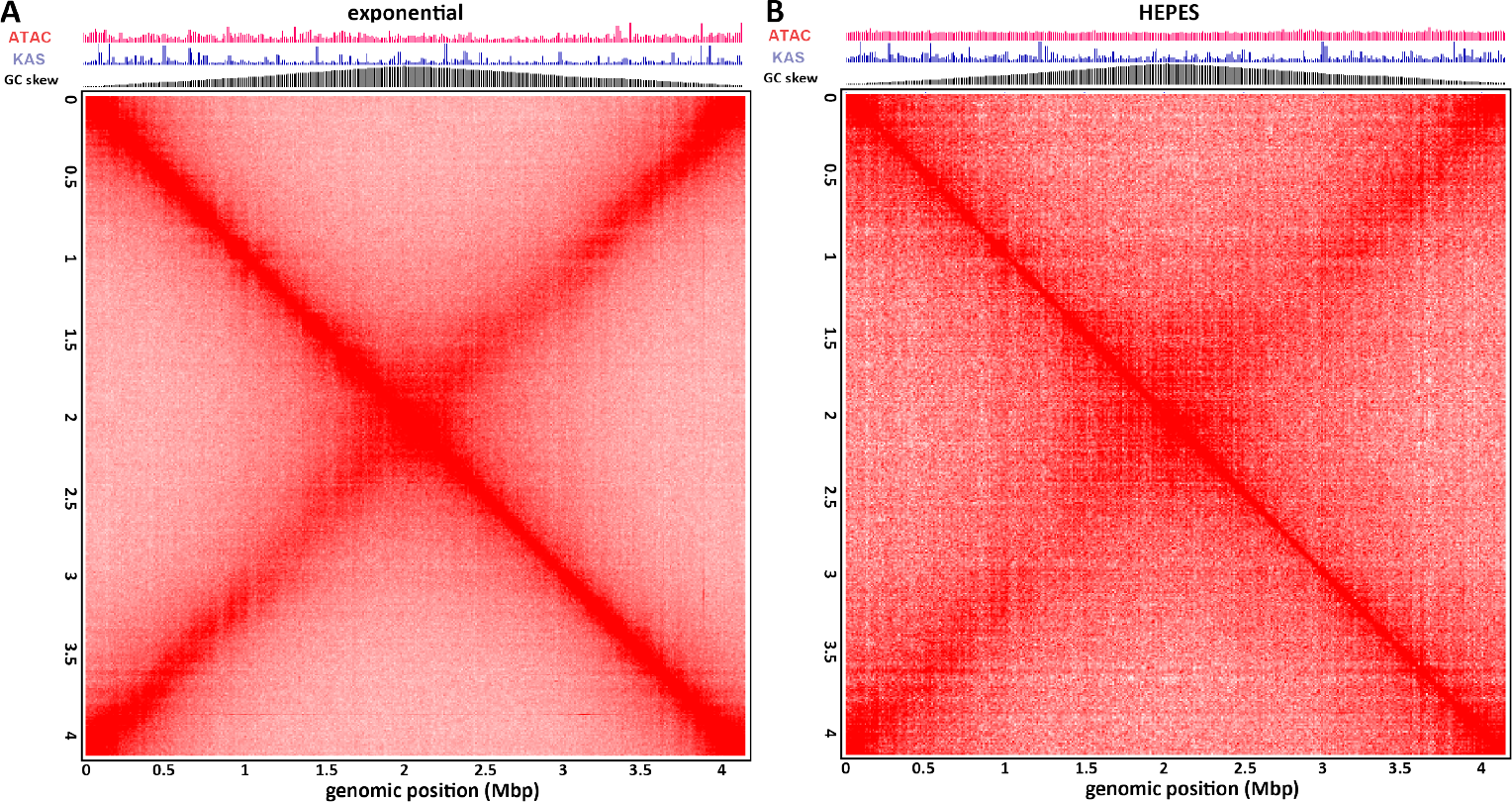
Three-dimension organization of the the *Bacteriovorax* sp. ICPB 3264 [H-I A3.12] chromosome. (A) Hi-C map of exponentially growing *Bacteriovorax* cells. (B) Hi-C map of starved (HEPES) *Bacteriovorax* cells.

## Discussion

The existence of bacteria with histone-based chromatin represents one of the most surprising and exciting discoveries in chromatin biology in recent years, coming after the assumption that histones are restricted to archaea and eukaryotes had reigned for many decades. Our study, carried out in a novel *Bacteriovorax* sp. strain whose genome we assembled and annotated *de novo*, presents the first direct measurements of chromatin accessibility, the distribution of active transcription and ssDNA, and three-dimensional genome conformation in such bacteria, and it allows us to chart some of the broadest trends in chromatin organization across the tree of life (Figure 7).

**Figure 7:**
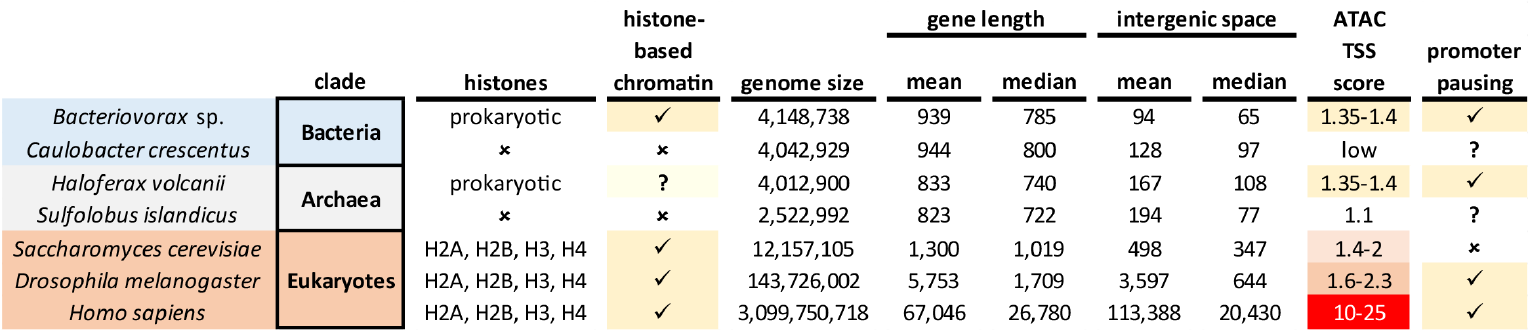
Comparison of the chromatin landscape across the deep organismal diversity. Shown is the presence of histones, histone-based chromatin, the basic genomic properties, i.e. genome size, gene length, gene density/intergenic space size, and the extent of concentration of ATAC-seq signal around promoters (“TSS score”, see the Methods section for details).

Like the euryarchaeon *Haloferax*, and similar to eukaryotes with nucleosomal chromatin, *Bacteriovorax* ‘s chromatin is characterized by preferentially accessible promoter regions. The relative level of promoter accessibility is comparable to that of *Haloferax*, slightly lower than yeast, and considerably lower than metazoans with much larger genomes. Likely this is a reflection of high transcriptional activity in extremely compacted genomes, i.e. actively transcribed regions temporarily lose the protection from transposase insertion conferred by histones or other packaging proteins, and the density of actively engaged polymerase molecules is higher in such cases than it is in mammals with highly bloated genomes. We see some more direct evidence for this phenomenon in the case of the rDNA operon in *Bacteriovorax*, which is the most broadly accessible gene in the genome, similar to the situation with rDNA in yeast.

Common with eukaryotes, but unlike the euryarchaeon *Haloferax, Bacteriovorax* exhibits positive correlation between chromatin accessibility and active transcription, suggesting that displacing histones from promoters might play a regulatory role in this organism. However, two observations complicate such an interpretation. First, we do not observe a protection signature in the ATAC-seq fragment length distribution analogous to what is seen in eukaryotes and also in some archaea with histone-based chromatin ^86^; it is thus not entirely clear how exactly *Bacteriovorax* histones physically protect DNA and bind to it (somewhat different models have been proposed in the past year ^26,27^). Second, in starved cells we observe loss of ATAC-seq enrichment over promoters but not of actively transcribing polymerases. This observation can be interpreted in two ways – either promoters close upon strong stress signals, implying that *Bacteriovorax* histones are not strongly inhibitory to polymerase activity, or, alternatively, histones are lost from chromatin, and this loss does not strongly affect polymerase engagement with DNA, even though histones appear to be essential in these organisms. Single-molecule mapping of chromatin accessibility ^33,34^ and examination of a much broader array of conditions can be expect to shed light on these questions.

Surprisingly, we also observe evidence for very strong polymerase pausing over *Bacteriovorax* genes, in contrast to what has been reported for bacteria such as *E. coli* ^53^, and similar to what is seen in many (though not all) eukaryotes. Regulation of productive transcriptional elongation might therefore be a key mechanism of modulating gene expression in bacteria with histone-based chromatin.

We also find evidence for independent regulation of and transcription initiation from internal promoters inside *Bacteriovorax* operons, similar to what is observed in the euryarchaeon *Haloferax*.

Finally, the *Bacteriovorax* genome exhibits three-dimensional organization typical for bacteria, centered on the axis defined by the replication origin and terminus. However, it does not display clear plectonemic CIDs. This interesting observation might further illuminate the mechanisms of formation or lack of such domains. Strong plectonemic CIDs are rare in eukaryotes with one notable exception – dinoflagellates, which display the largest and most pronounced such domains of any organism assayed so far, likely thanks to their unique genomic organization featuring unidirectional gene arrays many hundreds of kilobases long (thus generating extreme levels of supercoiling stress) and the also unique to them loss of histones as a main packaging component; it has been thus hypothesized that the frequent and strong interactions between nucleosomes in other eukaryotes overrides the formation of plectonemes driven by transcription-induced supercoiling, but in dinoflagellates that effect is unmasked ^87^. Conversely, if analogous such interactions are introduced in groups that normally do not feature histones and exhibit supercoiling domains, such as bacteria, it may be expected for CID-like domains to disappear. We do not observe clear CIDs in *Bacteriovorax*, but they are also not always robust in all bacteria, and not seen in the non-histone carrying crenarcheaote *Sulfolobus* while having been reported from *Haloferax*. However, more recent evidence points to *Haloferax* possessing histone genes but not actually having histone-based chromatin ^88,88–90^, in which case some other chromatin protein must be responsible for the high protection from Tn5 insertion measured over its genome, and other chromatin proteins might also be actively shaping the physical genome of other archaeons and bacteria. Further expanding the coverage of physical genome mapping over the deep diversity in the tree of life can be expected to resolve these and many other questions.

## Methods

Except where explicitly indicated otherwise, data was processed using custom-written Python scripts (https://github.com/georgimarinov/GeorgiScripts)

### Cell culture

We obtained the strain designated as *Bdellovibrio bacteriovorus* “ICPB 3264 [H-I A3.12]” from ATCC (https://www.atcc.org/products/25637). As discussed in the main text, subsequent analysis revealed that this strain does not in fact belong to the *Bdellovibrio bacteriovorus* clade, but is instead a *Bacteriovorax* strain that for which sequence data did not previously exist.

The cells were grown in ATCC medium 526 (10.0 g Peptone, 3.0 g Yeast extract, 1.0 L distilled water, autoclaved at 121 °C) at 30 °C, except where otherwise indicated for the HEPES starvation condition (in which case cells were centrifuged down and the media was replaced with HEPES buffer supplemented with 2 mM CaCl_2_ and 4 mM MgCl_2_ as previously described ^62^).

### Genome sequencing

Genomic DNA (gDNA) was isolated using the NEB Monarch Genomic DNA Purification Kit (Cat # T3010).

For Illumina sequencing, genomic DNA libraries were generated using Tn5 transposition as previously described ^31^. Briefly, ∼200 ng DNA gDNA were used, with the volume increased to 22.5 *μ*L using ultrapure H_2_O. Transposition was carried out with 22.5 *μ*L gDNA, 25 *μ*L 2 *×* TD buffer (20 mM Tris-HCl pH 7.6, 10 mM MgCl_2_, 20% Dimethyl Formamide), and 2.5 *μ*L Tn5, incubating at 37 °C for 15 minutes. Transposed DNA was immediately purified using the Qiagen MinElute PCR Purification Kit (Cat # 28004), with the reaction stopped with 250 *μ*L PB buffer, and elution in 10 *μ*L EB buffer. Final library amplification was carried out by mixing the 10 *μ*L eluate, 10 *μ*L H_2_O, 2.5 *μ*L i5 primer, 2.5 *μ*L i7 primer, and 25 *μ*L NEBNext High-Fidelity 2 *×* PCR Master Mix, using the following thermocycler program: 3 minutes at 72 °C, 30 seconds at 98 °C, 10 cycles of: 98 °C for 10 seconds, 63 °C for 30 seconds, 72 °C for 30 seconds. Final libraries were purified using the MinElute PCR Purification Kit.

Illumina libraries were sequenced in a 2 *×* 150 bp format using the Novogene sequencing service.

Nanopore sequencing was carried out using Flongle flowcells and the SQK-LSK114 library generation kit following the manufacturer’s instructions.

### Genome assembly

The `guppy` base caller (version 6.4.6) was used for nanopore base calling with the following settings: `--flowcell FLO-FLG114 --kit SQK-LSK114 –recursive --trim_strategy none`.

Genome assembly was generated using the SPAdes assembler ^91^ (version 3.15.4) in a hybrid nanopore/Illumina mode.

### Genome annotation

Genome annotation was carried out using the RAST web server ^92^. Ribosomal RNAs and other ncRNAs were annotated using Infernal ^93^ (version 1.1.1) and the RFAM database ^94^ (version 12.0) and RNAmmer ^95^ (version 1.2).

### Sequence analysis

DNA/RNA and protein multiple sequence alignment was carried out using ^96^.

Ribosomal RNA phylogenetic trees were built using RAxML-NG ^4^ (version 1.2.0) with the following settings: `--model GTR+G`, with all full-length or near full-length 16S rRNA sequences for Bdellovibrionota strains available on GenBank.

Protein sequences were translated from the newly generated genome annotation and available genome annotations for other sequenced Bdellovibrionota strains, and then protein domains were annotated using HMMER3^97^ (version 3.2.1) and the PFAM database ^98^ (version 32.0)

### Short-read 5-methylcytosine profiling

Genomic DNA was sheared on a Covaris E220, then converted into sequencing libraries following the EM-seq protocol, using the NEBNext Enzymatic Methyl-seq Kit (NEB, Cat # E7120L), and sequenced as 2 × 150mers on a NovaSeq X through Novogene.

Adapters were trimmed from sequencing reads using Trimmomatic ^101^ (version 0.36). Trimmed reads were aligned against our *de novo* genome assembly using `bwa-meth` with default settings. Duplicate reads were removed using `picard-tools` (version 1.99). Methylation calls were extracted using `MethylDackel` (https://github.com/dpryan79/MethylDackel). Additional analyses were carried out using custom-written Python scripts (https://github.com/georgimarinov/GeorgiScripts).

### ATAC-seq experiments

ATAC-seq experiments were carried out as previously described ^31,42^.

Cells were fixed by adding 37% formaldehyde (Sigma) at a final concentration of 1% and incubating for 15 minutes at room temperature. Formaldehyde was then quenched using 2.5 M glycine at a final concentration of 0.25 M. Cells were subsequently centrifuged at 13,000 rpm for 1 minute, washed once in 1 × PBS, and centrifuged again at 13,000 rpm for 1 minute. Lysis was then carried out by resuspending cells in 400 *μ*L Permeabilization Buffer (33 mM Tris-HCl pH 8.0, 20% sucrose, 25 *μ*g/mL lysozyme) and incubating 5 minutes at 30 °C in a ThermoMixer (with shaking at 1000 rpm). Cells were again pelleted at 13,000 rpm for 1 minute, resuspended in 50 *μ*L transposition mix (25 *μ*L 2*×* TD buffer, 2.5 *μ*L Tn5, 22.5 *μ*L ultrapure H_2_O), and incubated at 37 °C for 15 minutes. The reaction was stopped with the addition of 150 *μ*L IP Elution Buffer (1% SDS, 0.1 M NaHCO_3_) and 2 *μ*L Proteinase K (Promega, Cat # MC5005), then incubated at 65 °C overnight to reverse crosslinks. DNA was isolated by adding an equal volume of 25:24:1 phenol:chloroform:isoamyl solution, vortexing and centrifuging for 3 minutes at 14,000 rpm, then purifying the top aqueous phase using the MinElute PCR Purification Kit, eluting in 10 *μ*L EB buffer. Libraries were generated as already described above.

### ATAC-seq data processing

Demultipexed FASTQ files were mapped to the *de novo Bacteriovorax* sp. genome assembly as 2×36mers using Bowtie ^99^ (version 1.0.1) with the following settings: `-v 2 -k 2 -m 1 --best --strata`. Duplicate reads were removed using `picard-tools` (version 1.99).

TSS scores were calculated as the ratio of ATAC signal in the region *±*100 bp around TSSs versus the ATAC signal of the 100-bp regions centered at the two points ±2 kbp of the TSS as previously described ^43^.

Peak calling was carried out using MACS2^100^ with the following settings: `-g 4150000 -f BAM --to-large` `--keep-dup all --nomodel`.

### KAS-seq experiments

KAS-seq experiments were carried out following the previously published protocol ^35^ with some modifications in the sequencing library generation part ^31^.

Briefly, a 500-mM N_3_-kethoxal solution was brought to 37 °C, then added to 1 mL of culture at a final concentration of 5 *μ*M. Cells were then incubated for 5 minutes at 30 °C in a ThermoMixer at 1000 rpm.

Cells were then pelleted by centrifugation at 13,000 rpm for 1 minute, resuspended in 100 *μ*L 1 × PBS buffer, and DNA was immediately isolated using the Monarch Genomic DNA Purification Kit (NEB, Cat # T3010S), with the modification that elution was carried out with 87.5 *μ*L 25 mM K_3_BO_3_ solution (pH 7.0).

Biotin was clicked onto kethoxal-modified guanines by mixing 87.5 *μ*L DNA, 2.5 *μ*L 20 mM DBCO-PEG4-biotin (Sigma, Cat # 760749; DMSO solution), and 10 *μ*L 10 *×* PBS, and incubating at 37 °C for 90 minutes.

DNA was isolated using AMPure XP beads and eluted in 130 *μ*L 25 mM K_3_BO_3_ (pH 7.0), then sheared on a Covaris E220 for 120 seconds down to ∼150-200 bp.

Libraries were built on beads using the NEBNext Ultra II DNA Library Prep kit (NEB, Cat # E7645L). Biotin pull down was initiated py pipetting 20 *μ*L Dynabeads MyOne Streptavidin T1 beads (ThermoFisher Scientific, Cat # 65306) into DNA lo-bind tubes. Beads were separated on magnet, resuspended in 200 *μ*L of 1 *×* TWB buffer (Tween Washing Buffer; 5 mM Tris-HCl pH 7.5; 0.5 mM EDTA; 1 M NaCl; 0.05% Tween 20), then separated on magnet again and resuspended in 300 *μ*L of 2 *×* BB (Binding Buffer; 10 mM Tris-HCl pH 7.5, 1 mM EDTA; 2 M NaCl). The DNA (130 *μ*L) was added together with 170 *μ*L 0.1 *×* TE buffer, and incubated at RT on rotator for ≥15 minutes. Beads were separated on magnet, resuspended in 200 *μ*L of 1 *×* TWB, and incubated at 55 °C in a Termomixer for 2 minutes with shaking at 1000 rpm. Beads were again separated on magnet and the 200-*μ*L 55 °C TWB wash step was repeated. Beads were separated on magnet and resuspended in 50 *μ*L 0.1*×* TE.

End repair was carried out by adding 7 *μ*L NEB End Repair Buffer and 3 *μ*L NEB End Repair Enzyme, incubating at 20 °C for 30 minutes, then at 65 °C for 30 minutes.

End repair was followed by adaptor ligation by adding 2.5 *μ*L NEB Adaptor, 1 *μ*L NEB Ligation Enhancer and 30 *μ*L NEB Ligation Mix, incubating at 20 °C for 20 minutes, then adding 3 *μ*L USER Enzyme and incubating at 37 °C for 15 minutes. Beads were separated on magnet, resuspended in 200 *μ*L of 1 *×* TWB, then incubated at 55 °C in a Thermomixer for 2 minutes with shaking at 1000 rpm. Subsequently beads were separated on magnet and resuspended in 100 *μ*L of 0.1 *×* TE, separated on magnet again, resuspended in 15 *μ*L of 0.1 *×* TE Buffer, and transfered to PCR tubes.

Beads were then incubated at 98 °C for 10 minutes, and libraries were amplified by adding 5 *μ*L of i5 primer, 5 *μ*L of i7 primer and 25 *μ*L of 2 *×* Q5 Hot Start Polymerase Mix, using the following PCR program: 30 seconds at 98 °C; 15 cycles of 98 °C for 10 seconds, 65 °C for 30 seconds, and 72 °C for 30 seconds; and a final extension at 72 °C for 5 minutes.

Beads were separated on magnet and the final libraries were purified from the supernatant using 50 *μ*L AMPure XP beads, eluting in 0.1*×* TE buffer.

### KAS-seq data processing

Demultipexed FASTQ files were mapped to the *de novo Bacteriovorax* sp. genome assembly as 2 *×* 36mers using Bowtie ^99^ with the following settings: `-v 2 -k 2 -m 1 --best --strata`. Duplicate reads were removed using `picard-tools` (version 1.99).

### Differential accessibility/KAS-seq analysis

The analysis of differential chromatin accessibility as measured using ATAC-seq or enriched for KAS-seq signal was carried out using DESeq2^102^. Read counts were calculated over promoters or gene bodies and used as input into DESeq2.

### Hi-C experiments

Hi-C experiments were carried out as previously described ^87,103^ with some modifications.

Cells were crosslinked using 37% formaldehyde (Sigma) at a final concentration of 1% for 15 minutes at room temperature. Formaldehyde was then quenched using 2.5 M Glycine at a final concentration of 0.25 M. Cells were centrifuged at 13,000 rpm for 2 minutes, washed once in 1 *×* PBS, and stored at -80 °C.

Denaturation was carried out by resuspended cells in 50 *μ*L of 0.5% SDS and incubating at 62 °C for 10 minutes. SDS was quenched by adding 145 *μ*L of H_2_O and 25 *μ*L of 10% Triton X-100 and incubating at 37 °C for 15 minutes.

Restriction digestion was carried out by adding 25 *μ*L of 10*×* CutSmart buffer and 100 U of the MluCI restriction enzyme (NEB #R0538) plus 100 U of the MboI enzyme (NEB #R0147), then incubating for ≥2 hours at 37 °C in a Thermomixer at 900 rpm. The reaction was then incubated at 62 °C for 20 minutes in order to inactivate the restriction enzymes.

Fragment ends were filled in by adding 37.5 *μ*L of 0.4 mM biotin-14-dATP (ThermoFisher Scientific, # 19524-016), 1.5 *μ*L each of 10 mM dCTP, dGTP and dTTP, and 8 *μ*L of 5U/*μ*L DNA Polymerase I Large (Klenow) Fragment (NEB #M0210). The reaction was the incubated at 37 °C in a Thermomixer at 900 rpm for 45 minutes.

Fragment end ligation was carried out by adding 663 *μ*L H_2_O, 120 *μ*L 10 *×* NEB T4 DNA ligase buffer (NEB B0202), 100 *μ*L of 10% Triton X-100, 12 *μ*L of 10 mg/mL Bovine Serum Albumin (100 *×* BSA, NEB), 5 *μ*L of 400 U/*μ*L T4 DNA Ligase (NEB #M0202), and incubating at room temperature for ≥4 hours with rotation.

Cells were then pelleted by centrifugation at 13,000 rpm for 5 minutes. The pellet was resuspended in 200 *μ*L Elution Buffer (1% SDS, 0.1 M NaHCO_3_), Proteinase K was added, and incubated at 65 °C overnight to reverse crosslinks.

After addition of 600 *μ*L 1 *×* TE buffer, DNA was sheared using a Covaris E220 instrument. DNA was then purified using the MinElute PCR Purification Kit (Qiagen #28006), with elution in a total volume of 300 *μ*L 1*×* EB buffer.

For streptavidin pulldown of biotin-labeled DNA, 150 *μ*L of 10 mg/mL Dynabeads MyOne Streptavidin T1 beads (Life Technologies, 65602) were separated on a magnetic stand, then washed with 180 *μ*L of 1 *×* TWB (Tween Washing Buffer; 5 mM Tris-HCl pH 7.5; 0.5 mM EDTA; 1 M NaCl; 0.05% Tween 20). Beads were then resuspended in 300 *μ*L of 2 *×* Binding Buffer (10 mM Tris-HCl pH 7.5, 1 mM EDTA; 2 M NaCl), the sonicated DNA was added, and beads were incubated for ≥15 minutes at room temperature on a rotator. After separation on a magnetic stand, the beads were twice washed with 180 *μ*L of 1 *×* TWB, and heated at 55 °C in a Thermomixer with shaking for 2 minutes.

Final libraries were prepared on beads using the NEB-Next Ultra II DNA Library Prep Kit (NEB #E7645). End repair was carried out by resuspending beads in 50 *μ*L 1 *×* EB buffer, and adding 3 *μ*L NEB Ultra End Repair Enzyme and 7 *μ*L NEB Ultra End Repair Enzyme, followed by incubation at 20 °C for 30 minutes and then at 65 °C for 30 minutes.

Adapters were ligated to DNA fragments by adding 30 *μ*L Blunt Ligation mix, 1 *μ*L Ligation Enhancer and 2.5 *μ*L NEB Adapter, incubating at 20 °C for 20 minutes, adding 3 *μ*L USER enzyme, and incubating at 37 °C for 15 minutes. Beads were then separated on a magnetic stand, and washed with 180 *μ*L TWB for 2 minutes at 55 °C at 1000 rpm in a Thermomixer. After separation on a magnetic stand, beads were washed in 100 *μ*L 0.1 *×* TE buffer, then resuspended in 16 *μ*L 0.1 *×* TE buffer, and heated at 98 °C for 10 minutes.

For PCR, 5 *μ*L of each of the i5 and i7 NEB Next sequencing adapters were added together with 25 *μ*L 2 *×* NEB Ultra PCR Mater Mix. PCR was carried out with a 98 °C incubation for 30 seconds and 12 cycles of 98 °C for 10 seconds, 65 °C for 30 seconds, and 72 °C for 1 minute, followed by incubation at 72 °C for 5 minutes.

Beads were separated on a magnetic stand, and the supernatant was cleaned up using 1.8*×* AMPure XP beads.

Libraries were sequenced as 2 *×* 150mers on a Illumina NovaSeq X through Novogene.

### Hi-C data processing

Hi-C sequencing reads were processed against the *de novo Bacteriovorax* sp. assembly using the Juicer pipeline for analyzing Hi-C datasets ^104^ (version 1.6 of Juicer and version 2.13.07 of Juicer Tools) with default settings.

Hi-C matrices were visualized using Juicebox ^105^.

## Data availability

The sequencing datasets generated for and used in this study can be accessed from GEO accession GSE245010.

## Author contributions

G.K.M. conceptualized the study, carried out cell culture, and performed ATAC-seq, KAS-seq and Hi-C experiments together with B.D., analyzed the data and wrote the manuscript, with input from all authors. B.D. performed EM-seq experiments. W.J.G. and A.K. supervised the study.

## Acknowledgments

The authors would like to thank members of the Green-leaf and Kundaje labs for helpful discussions. This work was supported by NIH grants (P50HG007735, RO1 HG008140, U19AI057266 and UM1HG009442 to W.J.G., 1UM1HG009436 to W.J.G. and A.K., 1DP2OD022870-01 and 1U01HG009431 to A.K.), the Rita Allen Foundation (to W.J.G.), the Baxter Foundation Faculty Scholar Grant, and the Human Frontiers Science Program grant RGY006S (to W.J.G). W.J.G is a Chan Zuckerberg Biohub investigator and acknowledges grants 2017-174468 and 2018-182817 from the Chan Zuckerberg Initiative.

## Supplementary Materials

### Supplementary Figures

**Supplementary Figure 1:**
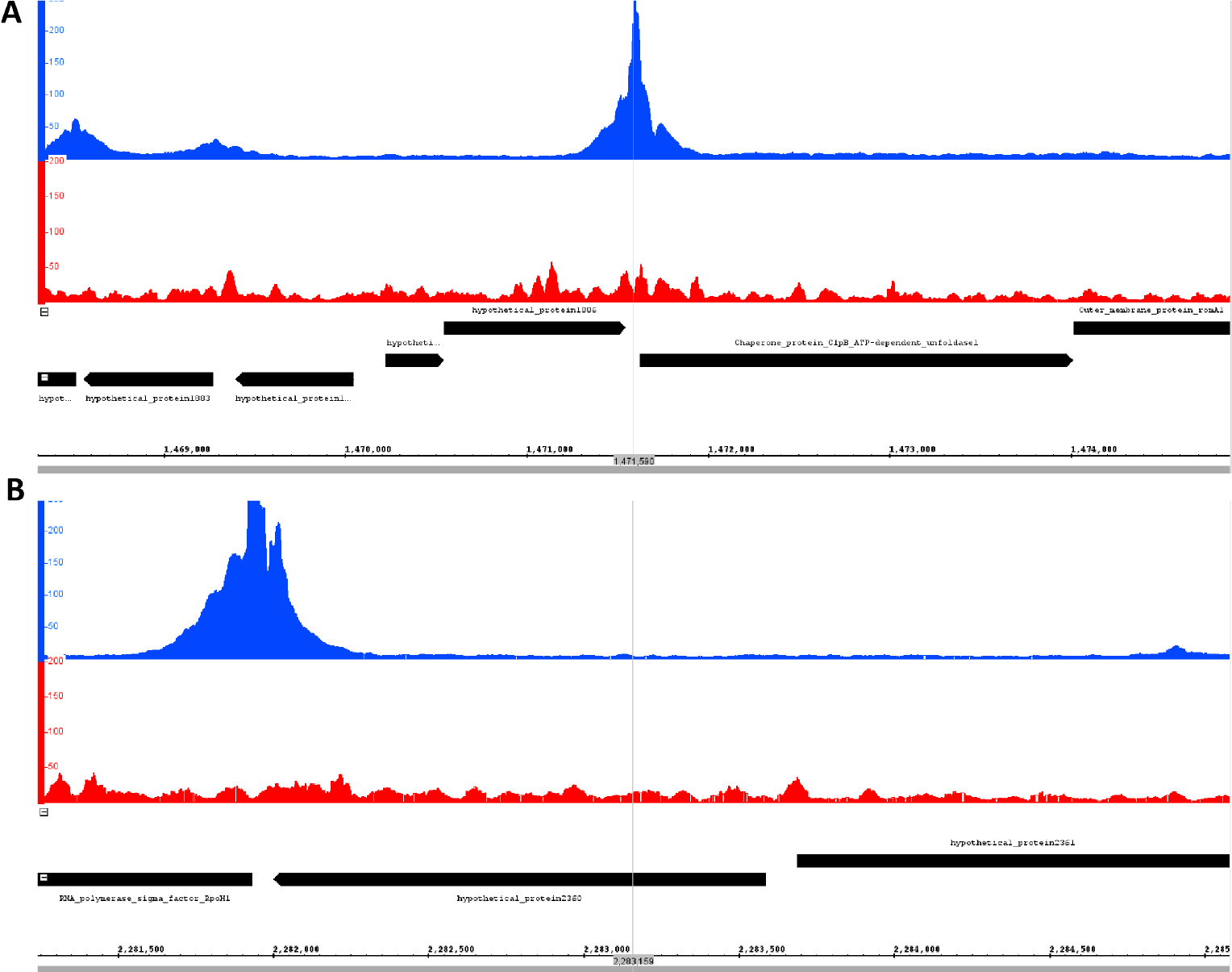
Additional examples of extremely strongly paused promoters in *Bacteriovorax*. (A) Chaperone protein ClpB ATP-dependent unfoldase (B) RNA Polymerase Sigma Factor RpoH1.

